# Detailed point cloud data on stem size and shape of Scots pine trees

**DOI:** 10.1101/2020.03.09.983973

**Authors:** Ninni Saarinen, Ville Kankare, Tuomas Yrttimaa, Niko Viljanen, Eija Honkavaara, Markus Holopainen, Juha Hyyppä, Saija Huuskonen, Jari Hynynen, Mikko Vastaranta

## Abstract

Quantitative assessment of the effects of forest management on tree size and shape has been challenging as there has been a lack of methodologies for characterizing differences and possible changes comprehensively in space and time. Terrestrial laser scanning (TLS) and photogrammetric point clouds provide three-dimensional (3D) information on tree stem reconstructions required for characterizing differences between stem shapes and growth allocation. This data set includes 3D reconstructions of stems of Scots pine (*Pinus sylvestris* L.) trees from sample plots with different thinning treatments. The thinning treatments include two intensities of thinning, three thinning types as well as control (i.e. no thinning treatment since the establishment). The data set can be used in developing point clouds processing algorithms for single tree stem reconstruction and for investigating variation in stem size and shape of Scots pine trees. Additionally, it offers possibilities in characterizing the effects of various thinning treatments on stem size and shape of Scots pine trees from boreal forests.

**Data set:** Zenodo https://zenodo.org/record/3701271

**Data set license:** Attribution 4.0 International (CC BY 4.0)

## Background and summary

In thinning, as a part of forest management, competition within a population is regulated as part of the trees are removed. Therefore, ecologically thinning is aimed at improving growing conditions (i.e., light, temperature, water, nutrients) of remaining trees and economically to maximize the net present value of a stand by decreasing the opportunity costs of the capital. Growing conditions are, thus, dependent on tree density that affects growth and structure of trees (Harper 1977) whereas changes in height and diameter growth alter stem form and taper. However, studies concentrating on thinning effects have only been based on a limited number of observations and the observations used have only been able to capture limited changes along a stem. Observations have included diameter-at-breast height (DBH), diameter at 6-m height, and tree height measured with calipers and clinometers. Differences in stem volume have been predicted using allometric equations that are incapable of considering variation in stem form. Thus, our understanding how forest management affects stem form and volume allocation is still rather limited.

Terrestrial laser scanning (TLS) has proven to non-destructively provide three-dimensional (3D) information on tree stems (Liang et al. 2014, Kankare et al. 2013, Raumonen et al. 2013, Saarinen et al. 2017) that has been impractical to produce with traditional means such as calipers or measurement tape. In applications related to boreal forests, TLS has been utilized in quantifying stem growth and changes in stem taper (Luoma et al. 2019), reparametrizing an existing taper curve model (Saarinen et al. 2019), and assessing timber quality (Pyörälä et al. 2019).

Detailed and quantitative information on stem size and shape has not been available for studies investigating variation in tree architecture as well as the effects of forest management and resultant growing conditions. Thus, 3D point cloud data were acquired for investigating the effects of different thinning treatments on stem size and shape of Scots pine (*Pinus Sylvestris* L.) trees without the need for destructive sampling (Saarinen et al. 2020a). The data set (Saarinen et al. 2020b) includes point cloud data from Scots pine trees from sample plots with different thinning treatments from southern Finland. This data set can be used in developing point cloud processing algorithms for single tree stem reconstruction and for investigating variation in stem shape as well as the effects of various thinning treatments on stem size and shape of Scots pine trees grown in boreal forests.

## Materials and methods

### Study site and data acquisition

The research was conducted in southern Finland in three study sites established and maintained by Natural Resources Institute Finland (Luke). Study site 1 was established in 2005 whereas study sites 2 and 3 in 2006. The study sites are located in the same vegetation zone namely southern Boreal forest zone at relative flat terrain, the elevation is 135 m, 155 m, and 120 m above sea level and the temperature sum is 1195, 1130, and 1256 degree days in study site 1, 2, and 3, respectively. Each study site is characterized as mesic heath forest (i.e., Myrtillus forest site type according to Cajander (1909)) and includes nine rectangular sample plots with size varying from 1000 m^2^ to 1200 m^2^ resulting in total of 27 sample plots.

The experimental design of the study sites includes two levels of thinning intensity and three thinning types resulting in six different thinning treatments, namely i) moderate thinning from below, ii) moderate thinning from above, iii) moderate systematic thinning, iv) intensive thinning from below, v) intensive thinning from above, and vi) intensive systematic thinning, as well as a control treatment where no thinning has been carried out since the establishment. Moderate thinning refers to prevailing thinning guidelines applied in Finland (Rantala 2011) whereas intensive thinning corresponds 50% lower remaining basal area (m^2^/ha) than in the plots with moderate thinning intensity. Regarding thinning types, small and suppressed trees as well as unsound and damaged trees (e.g. crooked, forked) were first removed from plots where thinning from below or above was carried out. Furthermore, suppressed and co-dominant trees were removed in thinning from below whereas mostly dominant trees were removed in thinning from above, but also maintaining regular spatial distribution of trees. In systematic thinning, on the other hand, only dominant trees were removed and small, suppressed trees were left to grow and regularity of spatial distribution of remaining trees was not emphasized similarly to other thinning types, although large gaps were avoided. Distribution of thinning treatments to sample plots within the three study sites is presented in Table 1.

**Table 1.** Thinning treatment of the sample plots in the three study sites.

| **Study site** | **Moderate below** | **Moderate above** | **Moderate systematic** | **Intensive below** | **Intensive above** | **Intensive systematic** | **No treatment** |
| --- | --- | --- | --- | --- | --- | --- | --- |
| **1** | 8 | 4, 9 | 5 | 7 | 1, 2 | 6 | 3 |
| **2** | 2 | 7 | 1, 8 | 5 | 6 | 4, 9 | 3 |
| **3** | 5 | 4 | 2, 8 | 3 | 6 | 1, 9 | 7 |

TLS data acquisition was carried out with Trimble TX5 3D laser scanner (Trible Navigation Limited, USA) for all three study sites between September and October 2018. Eight scans were placed to each sample plot, the scan setup is depicted in Figure 1, and scan resolution of point distance approximately 6.3 mm at 10-m distance was used. Artificial constant sized spheres (i.e. diameter of 198 mm) were placed around sample plots and used as reference objects for registering the eight scans onto a single, aligned coordinate system. The registration was carried out with FARO Scene software (version 2018).

**Figure 1.**
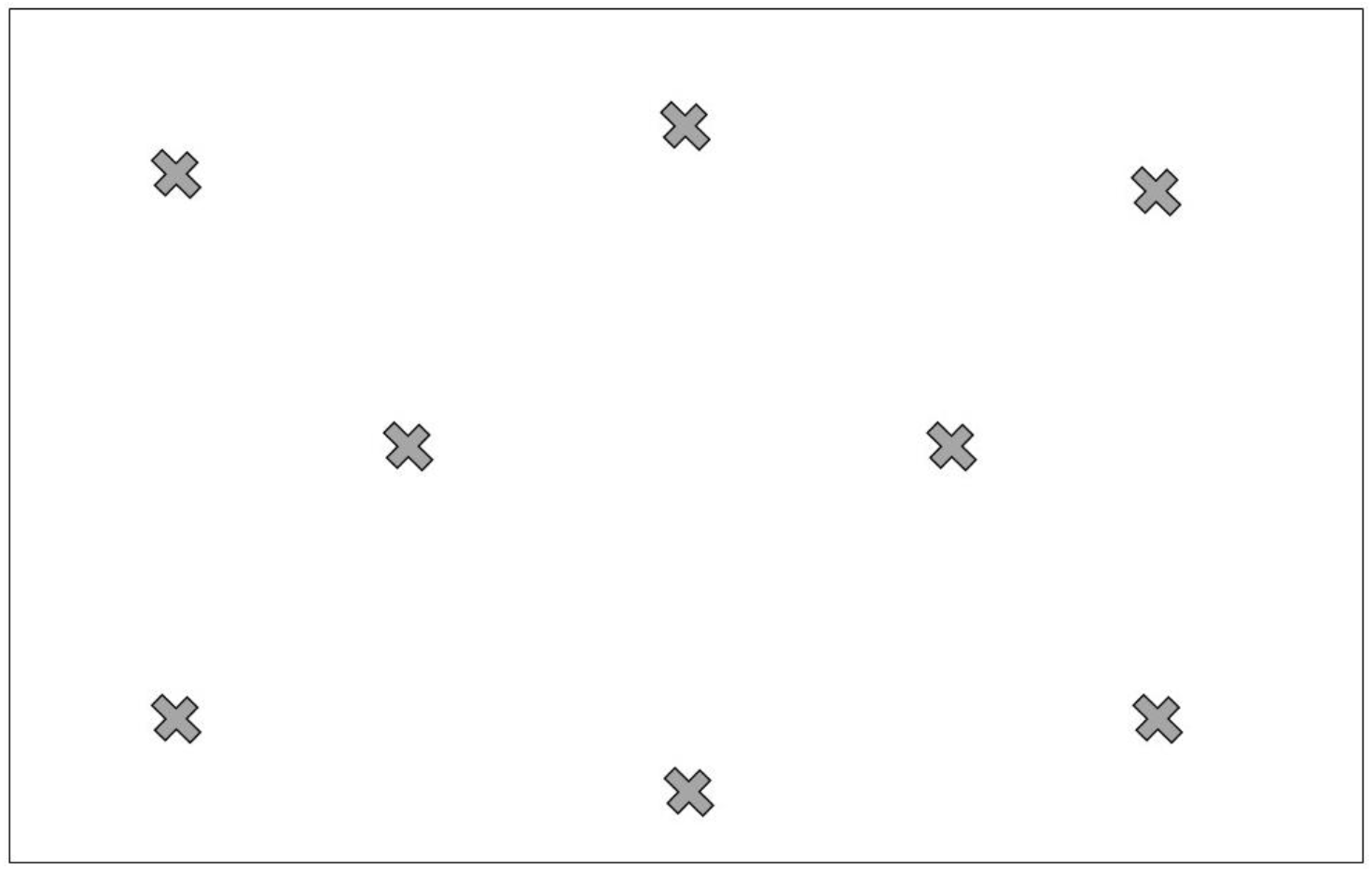
Scan design of eight scans (denoted as x) per an example sample plot.

In addition to TLS data, aerial images were obtained by using an unmanned aerial vehicle (UAV) with Gryphon Dynamics quadcopter frame. Two Sony A7R II digital cameras that had CMOS sensor of 42.4 MP, with a Sony FE 35mm f/2.8 ZA Carl Zeiss Sonnar T* lens, were mounted on the UAV in +15° and −15° angles. Images were acquired in every two seconds and image locations were recorded for each image. The flights were carried out on October 2, 2018. For each study site, eight ground control points (GCPs) were placed and measured. Flying height of 140 m and a flying speed of 5 m/s was selected for all the flights, resulting in 1.6 cm ground sampling distance. Total of 639, 614 and 663 images were captured for study site 1, 2, and 3, respectively, resulting in 93% and 75% forward and side overlaps, respectively. Photogrammetric processing of aerial images was carried out following the workflow as presented in Viljanen et al. (2018). The processing produced photogrammetric point clouds for each study site with point density of 804 points/m^2^, 976 points/m^2^, and 1030 points/m^2^ for study site 1, 2, and 3, respectively.

### Stem point extraction from TLS and UAV data

Point clouds from TLS and UAV were combined for characterizing Scots pine trees as comprehensively as possible. Individual tree detection and characterization from the combined point clouds followed the methodology presented by Yrttimaa et al. (2020) and Yrttimaa et al. (2019), respectively. The process included three steps: 1) point cloud normalization, 2) tree segmentation, and 3) point cloud classification. The TLS point clouds were normalized (i.e. point heights were transformed to heights above ground) using the *lasground* tool in LAStools software (Isenburg 2019) whereas RGB point clouds from UAV were normalized with digital terrain model (DTM) that is based on airborne laser scanning data and provided by National Land Survey of Finland (1). Canopy height models (CHMs) at a 20 cm resolution were generated from the normalized UAV point clouds to segment individual tree crowns. Variable Window Filter approach (Popescu & Wynne 2004) was used to identify the tree top positions, and Marker-Controlled Watershed Segmentation (Meyer & Beucher 1990) was applied to delineate the tree crown segments from the CHMs. The TLS point cloud was then split into tree-segments according to the extracted UAV crown segments using point-in-polygon approach (2). The tree-segmented point clouds were then classified into stem points and non-stem points. Points that represented planar, vertical, and cylindrical surfaces were classified as stem points while the rest of the points were classified as non-stem points (3).

## Data Records

This data set includes three packed zip files that can be downloaded from https://zenodo.org/record/3701271. The zip files include text files of stem points of each tree within the sample plots from the three test sites. The title of the zip file refers to the study sites 1, 2, and 3 described here. The title of the text files includes the information on the test site, the plot within the test site, and the tree within the plot. The text files contain stem points extracted from the point clouds.

The columns “x” and “y” contain x- and y-coordinates in a local coordinate system (in meters), in column “h” is the height of each point in meters above ground, and treeID is the tree identification number. The columns are separated by space. Based on the study site and plot number, files from different thinning treatments can be identified by using the information in Table 1.

## Technical Validation

At the sample plot level, TLS point clouds were co-registered with a mean distance error of 2.9 mm and standard deviation 1.2 mm, mean horizontal error was 1.3 mm (standard deviation 0.4 mm) and mean vertical error 2.3 mm (standard deviation 1.2 mm) (Saarinen et al. 2020) indicating high geometric accuracy of the point clouds. When similarly collected point clouds have been used to automatically measure tree diameters at multiple heights along a stem, root mean square error less than 1 cm can be expected in boreal forest conditions (Liang et al. 2018).

## Acknowledgements

The study was funded by the Academy of Finland, project numbers 315079, 272195, and 327861.

